# Characterization and treatment of SARS-CoV-2 in nasal and bronchial human airway epithelia

**DOI:** 10.1101/2020.03.31.017889

**Authors:** Andrés Pizzorno, Blandine Padey, Thomas Julien, Sophie Trouillet-Assant, Aurélien Traversier, Elisabeth Errazuriz-Cerda, Julien Fouret, Julia Dubois, Alexandre Gaymard, François-Xavier Lescure, Victoria Dulière, Pauline Brun, Samuel Constant, Julien Poissy, Bruno Lina, Yazdan Yazdanpanah, Olivier Terrier, Manuel Rosa-Calatrava

**Affiliations:** Virologie et Pathologie Humaine - VirPath team, Centre International de Recherche en Infectiologie (CIRI), INSERM U1111, CNRS UMR5308, ENS Lyon, Université Claude Bernard Lyon 1, Université de Lyon, Lyon, France; Signia Therapeutics SAS, Lyon, France; VirNext, Faculté de Médecine RTH Laennec, Université Claude Bernard Lyon 1, Université de Lyon, Lyon, France; Laboratoire Commun de Recherche HCL-bioMérieux, Centre Hospitalier Lyon-Sud, Pierre-Bénite, France; Centre d’Imagerie Quantitative Lyon-Est (CIQLE), Université Claude Bernard Lyon 1, Lyon, France; Laboratoire de Virologie, Centre National de Référence des virus Influenza Sud, Institut des Agents Infectieux, Groupement Hospitalier Nord, Hospices Civils de Lyon, Lyon, France; AP-HP, Infectious and Tropical Diseases Department, Bichat-Claude Bernard University Hospital, Paris, France; University of Paris, French Institute for Health and Medical Research (INSERM), IAME, U1137, Team DesCID, Paris, France; Epithelix Sàrl, Geneva, Switzerland; Pôle de Réanimation, Hôpital Roger Salengro, Centre Hospitalier Régional et Universitaire de Lille, Université de Lille 2, Lille, France

## Abstract

In the current COVID-19 pandemic context, proposing and validating effective treatments represents a major challenge. However, the lack of biologically relevant pre-clinical experimental models of SARS-CoV-2 infection as a complement of classic cell lines represents a major barrier for scientific and medical progress. Here, we advantageously used human reconstituted airway epithelial models of nasal or bronchial origin to characterize viral infection kinetics, tissue-level remodeling of the cellular ultrastructure and transcriptional immune signatures induced by SARS-CoV-2. Our results underline the relevance of this model for the preclinical evaluation of antiviral candidates. Foremost, we provide evidence on the antiviral efficacy of remdesivir and the therapeutic potential of the remdesivir-diltiazem combination as a rapidly available option to respond to the current unmet medical need imposed by COVID-19.

**One Sentence Summary:** New insights on SARS-CoV-2 biology and drug combination therapies against COVID-19.

## Main Text

On Dec 31, 2019, a cluster of cases of pneumonia of unknown etiology was reported in Wuhan, China. On Jan 10, 2020, a novel coronavirus, lately named severe acute respiratory syndrome coronavirus 2 (SARS-CoV-2) and classified into the Betacoronavirus genus, was identified as the causative agent (*1*). As of Mar 20, 2020, 9 days after the World Health Organization (WHO) declared the COVID-19 a pandemic (*2*), the new Coronavirus disease 2019 (COVID-19) had caused approximately 8 800 deaths among more than 210 000 confirmed cases reported mainly in China but also spreading to at least 168 other countries or territories worldwide (*3*). Compared to the two other coronaviruses responsible for epidemic outbreaks in the past, SARS-CoV and MERS-CoV, the novel SARS-CoV-2 strain shares ∼79% and ∼50% genome sequence identity, respectively (*4*). Not surprisingly, important differences in terms of the epidemiology and physiopathology between these three viruses have also been observed (*5–7*).

As with most emerging viral diseases, no specific antiviral treatment nor vaccine against any of these three coronaviruses are currently available, with standard patient management relying mainly on symptom treatment and respiratory support when needed. In that regard, and considering that key features of the biology of SARS-CoV-2 and its induced COVID-19 still require further characterization, the scarce readiness of biologically relevant pre-clinical experimental models of SARS-CoV-2 infection as a complement of the African green monkey VeroE6 cell line represents a major barrier for scientific and medical progress in this area. We and others have previously reported the advantage of using more physiological models such as in-house or commercially available reconstituted human airway epithelia (HAE) to isolate, culture and study a wide range of respiratory viruses (*8, 9*). Developed from biopsies of nasal or bronchial cells differentiated in the air/liquid interphase, these models reproduce with high fidelity most of the main structural, functional and innate immune features of the human respiratory epithelium that play a central role in the early stages of infection and constitute robust surrogates to study airway disease mechanisms and for drug discovery (*10*).

In this study, we initially isolated and amplified in VeroE6 cells a SARS-CoV-2 virus directly form a nasal swab from one of the first hospitalized patients with confirmed COVID-19 in France (*11*). The complete genome sequence of the isolated SARS-CoV-2 virus was deposited in the GISAID EpiCoVTM database under the reference BetaCoV/France/IDF0571/2020 (accession ID EPI_ISL_411218). Phylogenetic analysis confirmed that the isolated virus is representative of currently circulating strains (*12*). We first characterized the replicative capacities of this viral strain in VeroE6 cells at different multiplicities of infection (MOIs) (**Fig. 1A**), using both classic infectious titer determination in cell culture (TCID50) and molecular semi-quantitative methods, the latter based on ORF1b-nsp14-specific primers and probes designed by the School of Public Health/University of Hong Kong (details in Supplementary Materials). This double approach was facilitated by the appearance of clearly observable characteristic cytopathic effect from 48 hpi (**Fig. 1B**), and enabled the validation of a large interval (range 1-8 log10(TCID50)) with high correlation (R-squared 0.94) between molecular and infectious viral titers (**Fig. 1C**).

**Fig. 1.**
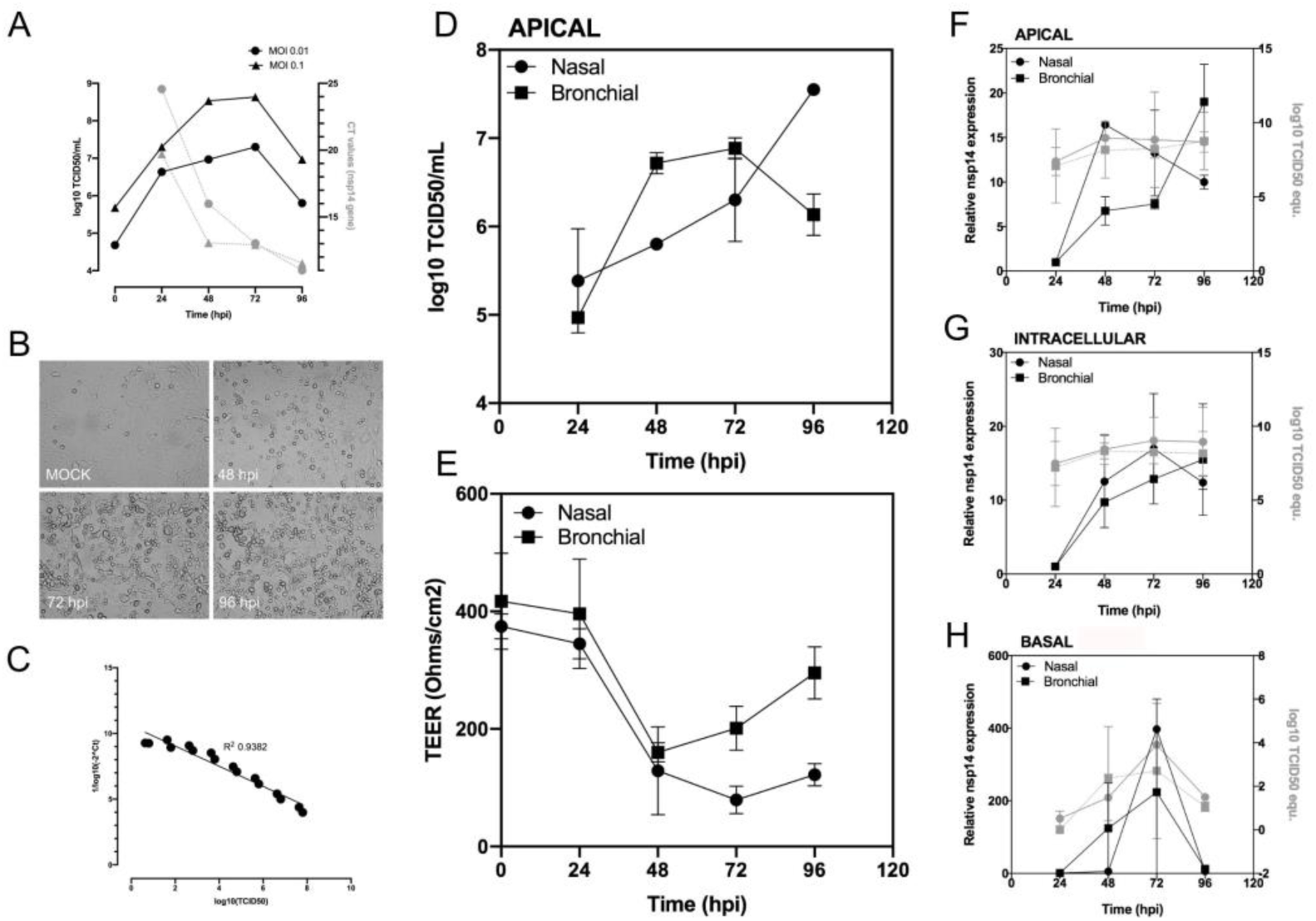
Characterization of a SARS-CoV-2 infection model in VeroE6 cells and in nasal and bronchial reconstituted human airway epithelia (HAE). **(A)** SARS-CoV-2 replication kinetics in VeroE6 cells. **(B)** Virus-induced cytopathic effects in VeroE6 cells. **(C)** Correlation between viral quantification methods. **(D)** Apical viral production was assessed in washes of the apical pole at 24, 48, 72 and 96 hpi. Vero E6 cells were incubated with serial dilutions of the collected sample for the determination of viral titers (log10 TCID50/ml) at the indicated time-points. **(E)** Trans-epithelial resistance (TEER in Ohms/cm^2^) between the apical and basal poles was measured at each time-point. **(F-H)** Relative viral genome quantification at the apical, intracellular and basal compartments of the HAE was performed after viral or total RNA extraction and RTqPCR. Results are expressed in fold change of nsp14 expression compared to 24 hpi and also as log10 TCID50 equivalent values. Representative data are shown from three independent experiments.

In parallel, we successfully inoculated nasal MucilAir™ HAE on the apical surface directly with nasal swab samples, as confirmed by transmission electron microscopy observations (**Fig. S1**). Characteristic features of coronavirus-induced cell ultrastructure remodeling were easily distinguishable in both the apical and basal sides of the HAE at 48 hpi, notably the high accumulation of progeny virions in mucus-producer goblet cells. Then, we advantageously exploited the MucilAir™ HAE model and in-house adapted protocols previously optimized for different respiratory viruses (*13*) to perform experimental infections with SARS-CoV-2. Viral replication was monitored through repeated sampling and TCID50 titration at the apical surface of HAE (**Fig. 1D**). Trans-epithelial electrical resistance (TEER), considered as a surrogate of epithelium integrity, was also measured during the time-course of infection (**Fig. 1E**). In parallel, comparative molecular viral genome quantification was performed at the three levels of the air/liquid HAE interphase: in apical washes (**Fig. 1F**, Apical), total cellular RNA (**Fig. 1G**, Intracellular) and basal medium (**Fig. 1H**, Basal). SARS-CoV-2 viral production at the epithelial apical surface increased sharply at 48 hpi, reaching 5.8 and 6.3 log10 TCID50/mL in nasal and bronchial HAE, respectively. The peak of viral replication was reached earlier in bronchial (48-72 hpi) than in nasal HAE, in which a progressive increase in infectious viral titers was observed until at least 96 hpi (**Fig. 1D**). This replication kinetics was validated by molecular viral genome quantification at the apical pole (**Fig. 1F**). High viral replication correlated with a reduction in epithelium integrity at 48 hpi, reflected by more than 2.8- and 4-fold decreases in bronchial and nasal HAE TEER values, respectively, followed by a partial recovery in the case of bronchial HAE (**Fig. 1E**). Moreover, viral production at the apical pole was well correlated with intracellular viral genome detection during infection, except for the nasal HAE at 48 hpi, in which a strong relative increase of nsp14 RNA was observed (**Fig. 1F**). Interestingly, viral genome was detected in the basal medium from 48 hpi, with the peak observed at 72 hpi (**Fig. 1H**) coinciding with the highest impact of SARS-CoV-2 infection on epithelium integrity.

To further characterize the biology of the SARS-CoV-2, we inoculated both nasal (**Fig. 2A, B**) and bronchial (**Fig. 2C, D**) HAE and analyzed the infection-induced remodeling of the cellular ultrastructure using transmission electron microscopy. At 48 hpi, both HAE exhibited a well-established infection, with ciliated, goblet and to a lesser extent basal cells showing active production of viral progeny. This observation is accordance with viral replication results described in **Fig. 1** and with a recent study reporting high expression levels of the SARS-CoV-2 cell receptor angiotensin-converting enzyme-2 (ACE2) in both ciliated and goblet respiratory cells (*14*). As previously observed in structural studies of other coronaviruses, notably SARS-CoV and MERS-CoV (*15–18*), we distinguished characteristic clusters in the perinuclear region of infected HAE cells. These clusters are mainly composed of numerous viral single-and double-membrane vesicles (DMV) and mitochondria (**Fig. 2A, A1, B, C, D**). Large electron-dense structures corresponding to the accumulation of viral material in active virus replication zones as well as typical double-membrane spherules containing pieces of membranes interspaced among virions being formed were also observed at 48 hpi (**Fig. 2B, D**). Moreover, double-membraned spherules containing numerous virions are noticeable near the plasmatic membranes (**Fig. 2B1, D2**). These spherules as well as several clusters of virions were observed mostly at the surface of ciliated cells (**Fig. 2A2**). These features are characteristic of the late stages of the viral cycle, hence confirming the capacity of the HAE to reproduce the asynchronous nature of infection.

**Fig. 2.**
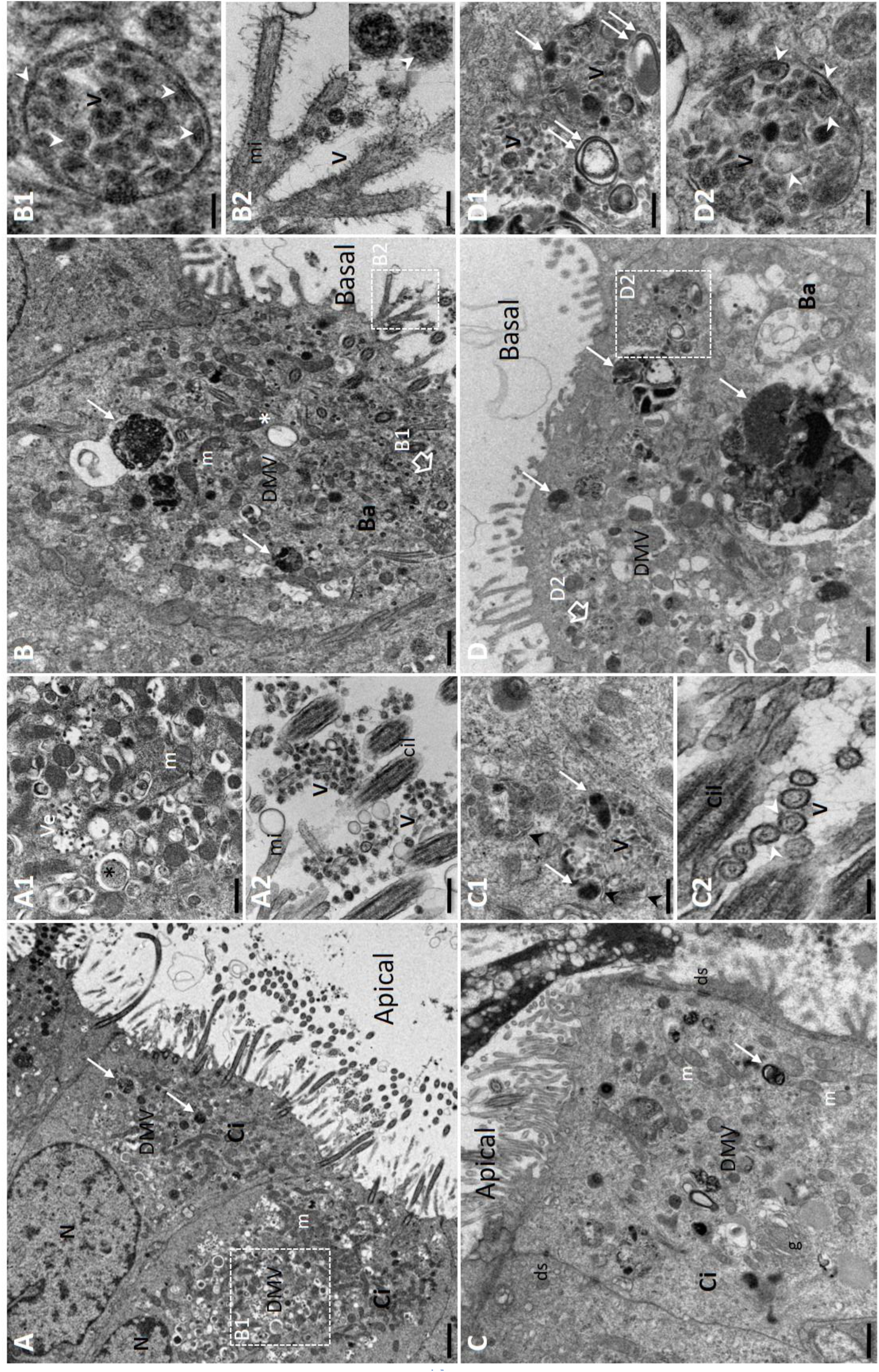
Ultrastructure of SARS-CoV-2 infected nasal and bronchial reconstituted human airway epithelia (HAE). MucilAir™ HAE were infected on the apical surface with SARS-CoV-2 (MOI 0.1). Forty-eight hours post inoculation, HAE were fixed and processed for transmission electron microscopy analysis, as described in Materials and Methods. **(A, B)** Section of apical ciliated (Ci) and basal (Ba) cells from nasal HAE showing numerous viral vesicles (DMVs) clustered in the perinuclear region in areas with mitochondria (m) and electron-dense accumulation of viral material (white arrow). Scale bar: 2 µm (A), 1 µm (B). **(A1)** Enlargement of cytoplasmic area with expended smooth-walled secretory vesicles containing virions (Ve) and virus-induced DMV (asterisk). Scale bar: 1 µm. **(A2)** Enlargement of ciliated cell surface showing virion clusters (V). Microvili (mi) and transverse sections of cilia (Cil) are observed. Scale bar: 0.05 µm. **(B1)** Enlargement of double-membraned spherules containing electron-dense material and pieces of double membrane interspaced among virions (V). Scale bar: 0.1 µm. **(B2)** Enlargement of Microvili (mi) and virons (V). White arrows point viral double membranes of virions seen at high magnification (inset). Scale bar: 0.2 µm. **(C, D)** Section of apical ciliated (Ci) and basal (Ba) cells from bronchial HAE showing numerous viral vesicles (DMVs) clustered in the perinuclear region in areas with mitochondria (m) and electron-dense accumulation of viral materials (white arrow). Scale bar: 1 µm. **(C1)** Enlargement of cytoplasmic area with spherules containing virions being formed (V), electron-dense accumulation of viral material (white arrow) and pieces of membranes (black arrowhead). Scale bar: 0.5 µm. **(C2)** High magnification of transverse section of virions (V) at the cell surface with cilia (Ci) and microvili (mi). Their double membrane (white arrowhead) and spikes at their outer edge are visible. Scale bar: 0.1 µm. **(D1)** Section of basal cells (Ba) showing virus-induced DMVs, large electron-dense accumulation of viral materials (white arrow) and double-membrane vesicles containing virions near the plasmatic membrane. Scale bar: 1 µm. **(E)** Enlargements of cytoplasmic area containing viral replication sites (double white arrows) and virions being formed (V). Scale bar: 0.1 µm. **(D2)** Enlargement of a double-membraned spherule containing virions (V), double-membrane vesicles and electron-dense viral materials. Scale bar: 0.1 µm. N: nucleus; DMV: cytoplasmic double-membrane vesicles; m: mitochondria; ds: desmosome.

Recent reports associate COVID-19 with high plasma levels of certain immunostimulant and proinflammatory cytokines (e.g. IL6), notably in patients with severe disease (*6*), suggesting a potential link with a poor prognosis (*19*). However, such inflammatory state has been much less characterized in the respiratory microenvironment thus far. To investigate the effect of SARS-CoV-2 infection on gene expression, we used Nanostring hybridization-based technology (Seattle, USA) on both nasal and bronchial HAE for multiplex mRNA detection and relative quantification of two complementary panels of genes (*20*) involved in the immune response (**Data S1**). Heatmap and hierarchical analyses identified two distinct levels of clustering. First, regardless of the nasal or bronchial nature of the HAE model, the gene expression profile at 24 hpi highlights a marked upregulation of a subset of ∼14% of the studied genes, which are subsequently downregulated at 48, 72 and 96 hpi (**Fig. 3A**). This observation was substantiated by unsupervised analysis of the data using the full gene panel. Indeed, the first component of the principal component analysis (PCA) that accounts for 63% of the variance is mainly driven by the time of infection, with a clear discrimination between 24 hpi and the other time-points (**Fig. 3B**, red triangles/dots). Interestingly, the immune transcriptomic signatures seem at least partially driven by the nature of the HAE beyond 24 hpi (**Fig. 3A**). This is in agreement with the second component analysis, gathering 12.6% of total variance, which allowed a clear differentiation between the nasal and bronchial compartments (**Fig. 3B**, green/purple triangles versus dots). In the nasal HAE, the response after 24 hpi (peak at 72-96 hpi) is driven by a strong upregulation of type I and type III IFNs (IFNB1, IFNL1, IFNL2-3-4) as well as other immunity-related genes. Whereas only a subset of these genes (CXCL10/IP10, CXCL2/MIP2A, IL1A, IL1B, Mx1 and ZBP1) follow the same pattern in the bronchial HAE, though at overall lower expression levels, the initial modulation of IFNB1, IFNL1, CCL2/MCP1 and IL6 in this tissue seems to fade at 96 hpi. Moreover, the relative expression of a subset of genes associated with the NF-kB and TNFα pathways (e.g. IL18, IL18R1, NFKB2, NFKBIA, TNFA, and TNFAIP3) is mostly unchanged all throughout the infection in bronchial HAE but it is highly upregulated in nasal HAE at 48 hpi and onwards (**Fig. 3C** and **Data S1**). Taken together, our results highlight distinctive transcriptional immune signatures between nasal and bronchial HAE, both in terms of kinetics and intensity, hence suggesting potential intrinsic differences in the early response to SARS-CoV-2 infection between the upper and lower respiratory tract. These results are in accordance with the first clinical reports describing in some patients a rapid worsening of the respiratory condition and overall clinical state by day 7-10 after symptom onset (*21*), most probably related to a cytokine storm syndrome (*22*).

**Fig. 3.**
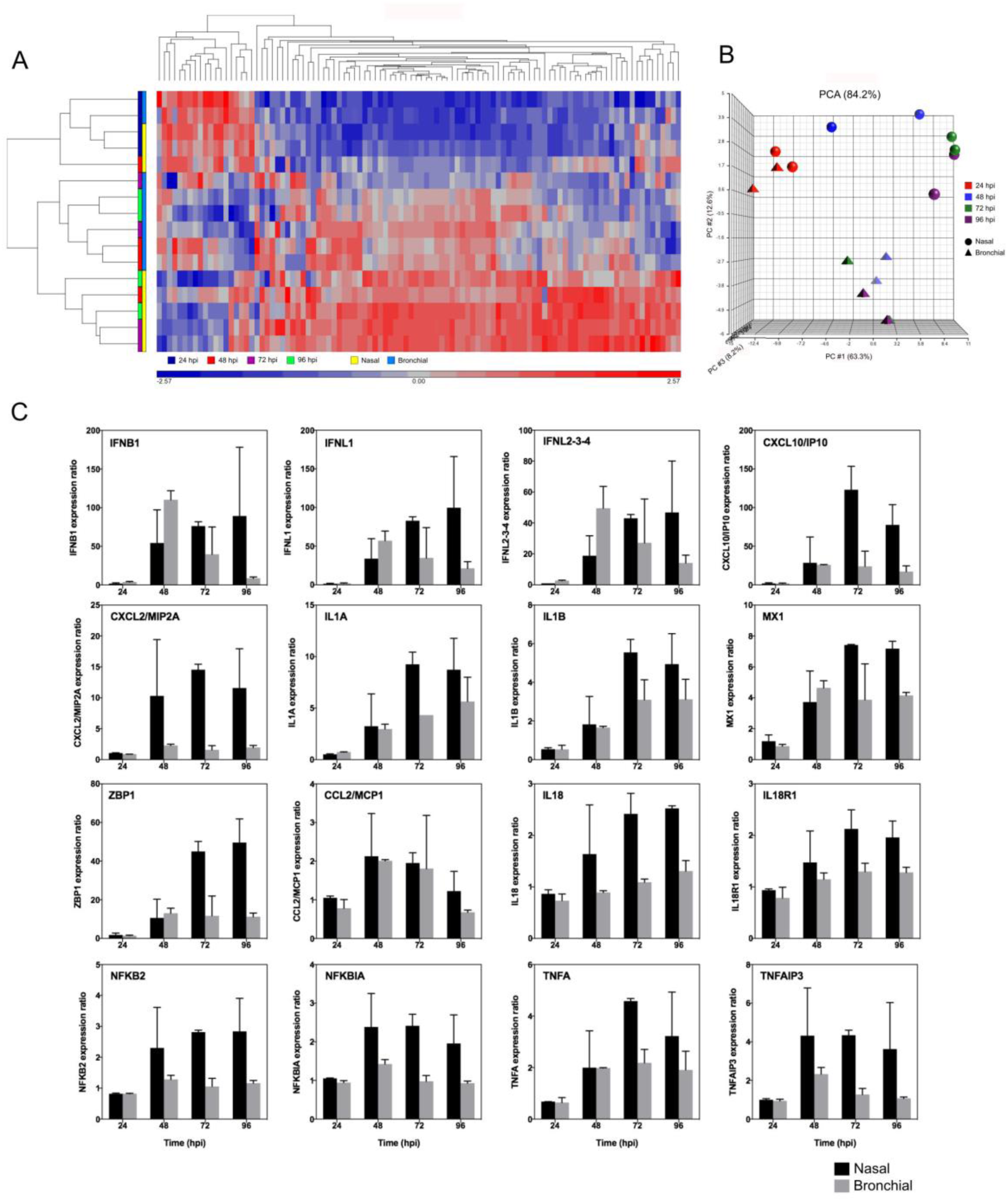
Nasal and bronchial innate immune transcriptional signature during the time course of SARS-COV-2 infection. Differential expression of both “immune response” (96 genes) and “type III IFNs” (12 genes) panels was evaluated in infected nasal and bronchial HAE using the Nanostring technology at the indicated time points. Data processing and normalization were performed with nSolver analysis software (version 4.0, NanoString technologies) and results are expressed in fold change induction compared to the mock condition. **(A)** Heatmap and hierarchical clustering of differentially expressed genes compared to the mock-infected condition. **(B)** Principal component analysis (PCA). **(C)** mRNA expression ratio of selected genes compared to the mock infected condition. Representative data are shown from two independent experiments.

The current absence of specific treatments against COVID-19 results in the empirical repurposing of approved or experimental drugs designed for other diseases. These treatments are usually based on limited clinical or preclinical data. Remdesivir (GS-5734) is a prodrug of an adenosine nucleotide analog with demonstrated broad antiviral activity against several RNA viruses in different preclinical models (*23*). Remdesivir has recently shown very promising results in animal models for the treatment of different coronaviruses, including MERS-CoV (*24*), as well as one *in vitro* study against SARS-CoV-2 (*25*). Indeed, various clinical trials on the use of remdesivir for the treatment of patients with COVID-19 have already started in China and USA. We therefore evaluated in both VeroE6 and HAE model the antiviral potential against SARS-CoV-2 of remdesivir monotherapy but also in combination with diltiazem. Diltiazem is a voltage gated Ca^2+^ channel antagonist currently used as anti-hypertensive for the control of angina pectoris and cardiac arrhythmia (*26*), which we have recently repurposed as an effective host-directed influenza inhibitor due to its so far undescribed capacity of inducing the interferon (IFN) antiviral response, particularly type III IFNs (**Fig. S2**) (*8*). Additionally, the rationale of testing such virus-directed plus host-directed drug combination is consistent with a novel study describing hypertension as a potential risk factor observed among a cohort of inpatients with COVID-19 (*19*), and two reports not anticipating potential adverse effects of diltiazem (*27*) or negative pharmacological interactions of between remdesivir and diltiazem for the treatment of COVID-19 (*28*).

**Figure 4A-C** shows a very strong antiviral effect of post-infection treatment with remdesivir in VeroE6 cells, with 50% inhibitory concentration (IC50) values of 0.98 ± 0.07 µM at 48 hpi and 0.72 ± 0.03 µM at 72 hpi. On the other hand, although the IFN-λ1 response in VeroE6 cells has been proved functional, this cell line cannot produce type I IFNs (*29, 30*). This incomplete IFN response most likely accounts for the lack of significant antiviral effect observed with diltiazem monotherapy in our experimental conditions. Nonetheless, addition of 11.5 µM diltiazem significantly potentiated the antiviral effect of remdesivir (**Fig. 4A-C**), inducing 68% and 50% reductions in remdesivir IC50 values at 48 and 72 hpi, respectively. Comparably, daily treatment with 20 µM remdesivir resulted in 7.3 log10 and 7.9 log10 reductions of intracellular SARS-CoV-2 viral titers at 48 hpi in nasal and bronchial HAE, respectively (**Fig. 4D**, upper panel). Not surprisingly for a model with a completely functional IFN response, daily treatment with 90 µM diltiazem resulted in moderate yet substantial (0.4 log10 and 0.8 log10, respectively) reductions of intracellular viral titers in nasal and bronchial HAE at the same time-point (**Fig. 4D**, upper panel). On top of that, we observed an additional 1.3 log10 reduction in nasal HAE viral titers for the remdesivir-diltiazem combination when compared with remdesivir monotherapy. In all cases, the antiviral effects induced by remdesivir, diltiazem or the remdesivir-diltiazem combination translated into a protection of the nasal but not the bronchial HAE barrier integrity, preventing the drop on TEER values induced by the infection (**Fig. 4D**, lower panel). Importantly, remdesivir also showed a strong antiviral effect at 72 hpi (**Fig. 4E**, upper panel). This time, the ∼2 log10 reductions in nasal and bronchial HAE viral titers observed for the remdesivir and remdesivir-diltiazem treatments correlated higher TEER values in both HAE compartments (**Fig. 4E**, lower panel).

**Fig. 4.**
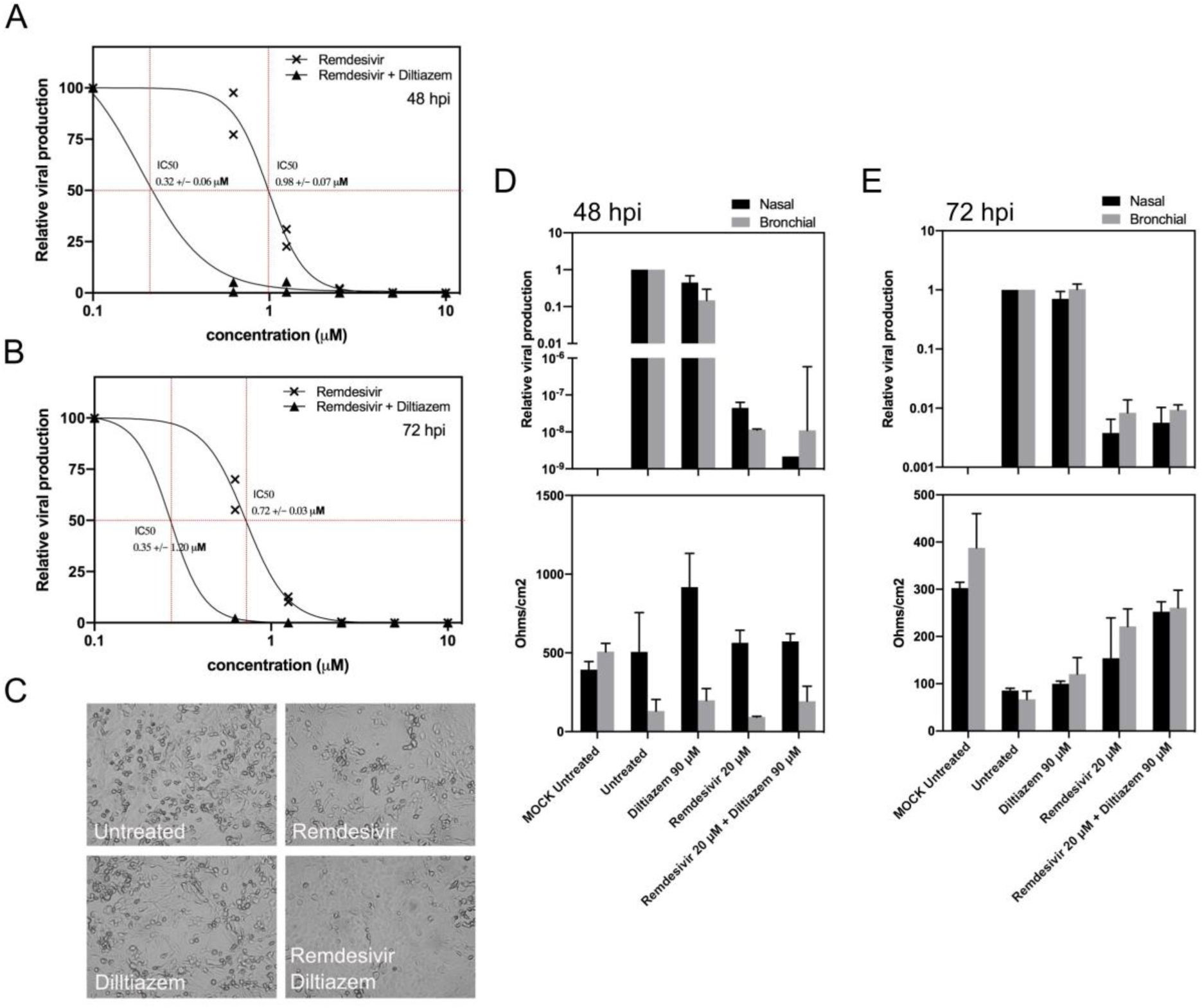
Diltiazem increases the antiviral activity of Remdesivir in VeroE6 cells and in HAE. **(A-B)** Dose-response curves of Remdesivir and Remdesivir-Diltiazem combination at 48 and 72 hpi in VeroE6 cells. **(C)** Effect of antiviral treatment on virally induced cytopathic effects in VeroE6 cells. **(D)** Relative intracellular viral genome quantification in nasal and bronchial HAE at 48 and 72 hpi. Results are expressed in relative viral production compared to the infected untreated control. **(D)** Trans-epithelial resistance (TEER in Ohms/cm2) between the apical and basal poles was measured at the same time points. Representative data are shown from three independent experiments.

Collectively, we report the utility of *in vitro* reconstituted HAE as an efficient support for the study of respiratory viral infections and virus-host interactions in highly biologically relevant experimental conditions. Our results obtained in this model provide novel contributions to the characterization of the viral infection and replication kinetics as well as on the tissue-level remodeling of the cellular ultrastructure induced by SARS-CoV-2. Interestingly, Nanostring results highlight a differential effect of SARS-CoV-2 infection on the early innate immune response of nasal and bronchial HAE. These responses are in line with the viral replication kinetics, and in some cases (*6*) -but not in others- (*31*) follow similar trends to what has been described in blood samples from patient cohorts. We expect our results will provide a starting point for future studies aimed at further characterizing the local pathophysiology and immune response to SARS-CoV-2 infection, particularly in the lower respiratory tract, with the ultimate objective of providing insight in terms of putative prognostic biomarkers and/or patient management.

Finally, the HAE model of SARS-CoV-2 infection described in this study also constitutes a very interesting physiologic model to evaluate candidate therapeutic approaches, provided that in many cases inhibitory effects observed in classic models of immortalized cell lines do not necessarily translate into the real clinical setting (*32*). In that regard, although further studies in terms of dose optimization would be desired, we provide evidence in both classic and complex *in vitro* experimental models on the potential of the remdesivir-diltiazem combination as an option worthy of consideration to respond to the current medical unmet need imposed by COVID-19. The combination of a virus-directed plus a host-directed drug could result in enhanced antiviral and/or immunomodulatory effects, including during the immunopathologic phase frequently observed the second week of infection. This association could also reduce the therapeutic doses of chemotherapeutic agents targeting nucleic acid synthesis and therefore minimize putative adverse side effects. Indeed, such two-pronged approach would be of special interest for the treatment of at-risk patients within the first 10 days from symptom onset, when effective clearance of viral load still plays a major role in the clinical outcome.

## Acknowledgments

The authors want to acknowledge Denis Ressnikoff and the CIQLE team (Université Claude Bernard Lyon 1) for their technical assistance with electron microscopy image acquisition and analysis as well as Dr. Guy Boivin for his critical reading of the manuscript.

## Author contributions

conceptualization: AP, OT, MRC. Formal analysis: AP, STA, JF, OT. Funding acquisition: BL, OT, MRC. Investigation: AP, BP, TJ, STA, AT, EEC, VD, JD, OT. Methodology: SC (MucilAir™ HAE). Project administration: OT, MRC. Resources: AG, SC, BL, YY. Supervision: AP, OT, MRC. Validation: BL, OT, MRC. Visualization: AP, STA, OT, MRC. Writing - original draft: AP, OT, MRC. Writing - review & editing: SC, JP, BL, YY, MRC.

## Data and materials availability

All data is available in the main text or the supplementary materials.

## Supplementary Materials

Materials and Methods

Figures S1-S2

Data S1

## References

1. D. S. Hui, E. I Azhar, T. A. Madani, F. Ntoumi, R. Kock, O. Dar, G. Ippolito, T. D. Mchugh, Z. A. Memish, C. Drosten, A. Zumla, E. Petersen, The continuing 2019-nCoV epidemic threat of novel coronaviruses to global health - The latest 2019 novel coronavirus outbreak in Wuhan, China. Int. J. Infect. Dis. 91, 264–266 (2020).

2. WHO Director-General’s opening remarks at the media briefing on COVID-19 - 11 March 2020, (available at https://www.who.int/dg/speeches/detail/who-director-general-s-opening-remarks-at-the-media-briefing-on-covid-19---11-march-2020).

3. nCoV 2019 situation (public), (available at https://who.maps.arcgis.com/apps/opsdashboard/index.html#/c88e37cfc43b4ed3baf977d77e4a0667).

4. R. Lu, X. Zhao, J. Li, P. Niu, B. Yang, H. Wu, W. Wang, H. Song, B. Huang, N. Zhu, Y. Bi, X. Ma, F. Zhan, L. Wang, T. Hu, H. Zhou, Z. Hu, W. Zhou, L. Zhao, J. Chen, Y. Meng, J. Wang, Y. Lin, J. Yuan, Z. Xie, J. Ma, W. J. Liu, D. Wang, W. Xu, E. C. Holmes, G. F. Gao, G. Wu, W. Chen, W. Shi, W. Tan, Genomic characterisation and epidemiology of 2019 novel coronavirus: implications for virus origins and receptor binding. Lancet. 395, 565–574 (2020).

5. M. Cascella, M. Rajnik, A. Cuomo, S. C. Dulebohn, R. Di Napoli, in StatPearls (StatPearls Publishing, Treasure Island (FL), 2020; http://www.ncbi.nlm.nih.gov/books/NBK554776/).

6. C. Huang, Y. Wang, X. Li, L. Ren, J. Zhao, Y. Hu, L. Zhang, G. Fan, J. Xu, X. Gu, Z. Cheng, T. Yu, J. Xia, Y. Wei, W. Wu, X. Xie, W. Yin, H. Li, M. Liu, Y. Xiao, H. Gao, L. Guo, J. Xie, G. Wang, R. Jiang, Z. Gao, Q. Jin, J. Wang, B. Cao, Clinical features of patients infected with 2019 novel coronavirus in Wuhan, China. Lancet. 395, 497–506 (2020).

7. Y. Wang, Y. Wang, Y. Chen, Q. Qin, Unique epidemiological and clinical features of the emerging 2019 novel coronavirus pneumonia (COVID-19) implicate special control measures. J. Med. Virol. (2020), doi:10.1002/jmv.25748.

8. A. Pizzorno, O. Terrier, C. Nicolas de Lamballerie, T. Julien, B. Padey, A. Traversier, M. Roche, M.-E. Hamelin, C. Rhéaume, S. Croze, V. Escuret, J. Poissy, B. Lina, C. Legras-Lachuer, J. Textoris, G. Boivin, M. Rosa-Calatrava, Repurposing of Drugs as Novel Influenza Inhibitors From Clinical Gene Expression Infection Signatures. Front Immunol. 10, 60 (2019).

9. S. M. Jazaeri Farsani, M. Deijs, R. Dijkman, R. Molenkamp, R. E. Jeeninga, M. Ieven, H. Goossens, L. van der Hoek, Culturing of respiratory viruses in well-differentiated pseudostratified human airway epithelium as a tool to detect unknown viruses. Influenza and Other Respiratory Viruses. 9, 51–57 (2015).

10. B. Boda, S. Benaoudia, S. Huang, R. Bonfante, L. Wiszniewski, E. D. Tseligka, C. Tapparel, S. Constant, Antiviral drug screening by assessing epithelial functions and innate immune responses in human 3D airway epithelium model. Antiviral Research. 156, 72–79 (2018).

11. F.-X. Lescure, L. Bouadma, D. Nguyen, M. Parisey, P.-H. Wicky, S. Behillil, A. Gaymard, M. Bouscambert-Duchamp, F. Donati, Q. Le Hingrat, V. Enouf, N. Houhou-Fidouh, M. Valette, A. Mailles, J.-C. Lucet, F. Mentre, X. Duval, D. Descamps, D. Malvy, J.-F. Timsit, B. Lina, S. Van Der Werf, Y. Yazdanpanah, Clinical and virological data of the first cases of COVID-19 in Europe: a case series. Lancet Infectious Diseases. In Press.

12. GISAID - Next hCoV-19 App, (available at https://www.gisaid.org/epiflu-applications/next-hcov-19-app/).

13. C. Nicolas de Lamballerie, A. Pizzorno, J. Dubois, T. Julien, B. Padey, M. Bouveret, A. Traversier, C. Legras-Lachuer, B. Lina, G. Boivin, O. Terrier, M. Rosa-Calatrava, Characterization of cellular transcriptomic signatures induced by different respiratory viruses in human reconstituted airway epithelia. Sci Rep. 9, 11493 (2019).

14. W. Sungnak, N. Huang, C. Bécavin, M. Berg, H. L. B. Network, SARS-CoV-2 Entry Genes Are Most Highly Expressed in Nasal Goblet and Ciliated Cells within Human Airways. 2003.06122 [q-bio] (2020) (available at http://arxiv.org/abs/2003.06122).

15. K. Knoops, M. Kikkert, S. H. E. van den Worm, J. C. Zevenhoven-Dobbe, Y. van der Meer, A. J. Koster, A. M. Mommaas, E. J. Snijder, SARS-coronavirus replication is supported by a reticulovesicular network of modified endoplasmic reticulum. PLoS Biol. 6, e226 (2008).

16. C. S. Goldsmith, K. M. Tatti, T. G. Ksiazek, P. E. Rollin, J. A. Comer, W. W. Lee, P. A. Rota, B. Bankamp, W. J. Bellini, S. R. Zaki, Ultrastructural characterization of SARS coronavirus. Emerging Infect. Dis. 10, 320–326 (2004).

17. B. A. Afzelius, Ultrastructure of human nasal epithelium during an episode of coronavirus infection. Virchows Arch. 424, 295–300 (1994).

18. A. H. de Wilde, V. S. Raj, D. Oudshoorn, T. M. Bestebroer, S. van Nieuwkoop, R. W. A. L. Limpens, C. C. Posthuma, Y. van der Meer, M. Bárcena, B. L. Haagmans, E. J. Snijder, B. G. van den Hoogen, MERS-coronavirus replication induces severe in vitro cytopathology and is strongly inhibited by cyclosporin A or interferon-α treatment. J. Gen. Virol. 94, 1749–1760 (2013).

19. F. Zhou, T. Yu, R. Du, G. Fan, Y. Liu, Z. Liu, J. Xiang, Y. Wang, B. Song, X. Gu, L. Guan, Y. Wei, H. Li, X. Wu, J. Xu, S. Tu, Y. Zhang, H. Chen, B. Cao, Clinical course and risk factors for mortality of adult inpatients with COVID-19 in Wuhan, China: a retrospective cohort study. Lancet (2020), doi:10.1016/S0140-6736(20)30566-3.

20. W. Mouton, C. Albert-Vega, M. Boccard, F. Bartolo, G. Oriol, J. Lopez, A. Pachot, J. Textoris, F. Mallet, K. Brengel-Pesce, S. Trouillet-Assant, Towards standardization of immune functional assays. Clin. Immunol. 210, 108312 (2020).

21. N. Zhu, D. Zhang, W. Wang, X. Li, B. Yang, J. Song, X. Zhao, B. Huang, W. Shi, R. Lu, P. Niu, F. Zhan, X. Ma, D. Wang, W. Xu, G. Wu, G. F. Gao, W. Tan, A Novel Coronavirus from Patients with Pneumonia in China, 2019. N Engl J Med. 382, 727–733 (2020).

22. P. Mehta, D. F. McAuley, M. Brown, E. Sanchez, R. S. Tattersall, J. J. Manson, COVID- 19: consider cytokine storm syndromes and immunosuppression. The Lancet, S0140673620306280 (2020).

23. T. P. Sheahan, A. C. Sims, R. L. Graham, V. D. Menachery, L. E. Gralinski, J. B. Case, S. R. Leist, K. Pyrc, J. Y. Feng, I. Trantcheva, R. Bannister, Y. Park, D. Babusis, M. O. Clarke, R. L. Mackman, J. E. Spahn, C. A. Palmiotti, D. Siegel, A. S. Ray, T. Cihlar, R. Jordan, M. R. Denison, R. S. Baric, Broad-spectrum antiviral GS-5734 inhibits both epidemic and zoonotic coronaviruses. Sci Transl Med. 9 (2017), doi:10.1126/scitranslmed.aal3653.

24. T. P. Sheahan, A. C. Sims, S. R. Leist, A. Schäfer, J. Won, A. J. Brown, S. A. Montgomery, A. Hogg, D. Babusis, M. O. Clarke, J. E. Spahn, L. Bauer, S. Sellers, D. Porter, J. Y. Feng, T. Cihlar, R. Jordan, M. R. Denison, R. S. Baric, Comparative therapeutic efficacy of remdesivir and combination lopinavir, ritonavir, and interferon beta against MERS-CoV. Nat Commun. 11, 222 (2020).

25. M. Wang, R. Cao, L. Zhang, X. Yang, J. Liu, M. Xu, Z. Shi, Z. Hu, W. Zhong, G. Xiao, Remdesivir and chloroquine effectively inhibit the recently emerged novel coronavirus (2019-nCoV) in vitro. Cell Res. 30, 269–271 (2020).

26. 51 PubChem, Diltiazem, (available at https://pubchem.ncbi.nlm.nih.gov/compound/39186).

27. L. Fang, G. Karakiulakis, M. Roth, Are patients with hypertension and diabetes mellitus at increased risk for COVID-19 infection? The Lancet Respiratory Medicine (2020), doi:10.1016/S2213-2600(20)30116-8.

28. Liverpool COVID-19 Interactions, (available at http://www.covid19-druginteractions.org/).

29. J. M. Emeny, M. J. Morgan, Regulation of the interferon system: evidence that Vero cells have a genetic defect in interferon production. J. Gen. Virol. 43, 247–252 (1979).

30. J. Prescott, P. Hall, M. Acuna-Retamar, C. Ye, M. G. Wathelet, H. Ebihara, H. Feldmann, B. Hjelle, New World Hantaviruses Activate IFNλ Production in Type I IFN-Deficient Vero E6 Cells. PLoS ONE. 5, e11159 (2010).

31. I. Thevarajan, T. H. O. Nguyen, M. Koutsakos, J. Druce, L. Caly, C. E. van de Sandt, X. Jia, S. Nicholson, M. Catton, B. Cowie, S. Y. C. Tong, S. R. Lewin, K. Kedzierska, Breadth of concomitant immune responses prior to patient recovery: a case report of non-severe COVID-19. Nature Medicine (2020), doi:10.1038/s41591-020-0819-2.

32. B. Cao, Y. Wang, D. Wen, W. Liu, J. Wang, G. Fan, L. Ruan, B. Song, Y. Cai, M. Wei, X. Li, J. Xia, N. Chen, J. Xiang, T. Yu, T. Bai, X. Xie, L. Zhang, C. Li, Y. Yuan, H. Chen, H. Li, H. Huang, S. Tu, F. Gong, Y. Liu, Y. Wei, C. Dong, F. Zhou, X. Gu, J. Xu, Z. Liu, Y. Zhang, H. Li, L. Shang, K. Wang, K. Li, X. Zhou, X. Dong, Z. Qu, S. Lu, X. Hu, S. Ruan, S. Luo, J. Wu, L. Peng, F. Cheng, L. Pan, J. Zou, C. Jia, J. Wang, X. Liu, S. Wang, X. Wu, Q. Ge, J. He, H. Zhan, F. Qiu, L. Guo, C. Huang, T. Jaki, F. G. Hayden, P. W. Horby, D. Zhang, C. Wang, A Trial of Lopinavir–Ritonavir in Adults Hospitalized with Severe Covid- New England Journal of Medicine (2020), doi:10.1056/NEJMoa2001282.

