## Supplementary files for "Characterization and treatment of SARS-CoV-2 in nasal and bronchial human airway epithelia"

**Affiliations:**

**This PDF file includes:**

Materials and Methods
Figs. S1 to S2
Data S1

### Materials and Methods

Clinical samples, viral isolation and sequencing

The SARS-CoV-2 strain used in this study was isolated by directly inoculating VeroE6 cell monolayers with a nasal swab sample collected from a one of the first COVID-19 cases confirmed in France: a 47y-o female patient hospitalized in January 2020 in the Department of Infectious and Tropical Diseases, Bichat Claude Bernard Hospital, Paris *(11)*. Once characteristic CPE was observable in more than 50% of the cell monolayer, supernatants were collected and immediately stored at -80 °C for subsequent viral RNA extraction using the QiAmp viral RNA Kit (Qiagen). The complete viral genome sequence was obtained using Illumina MiSeq sequencing technology, was then deposited after assembly on the GISAID EpiCoV platform (Accession ID EPI_ISL_411218) under the name BetaCoV/France/IDF0571/2020.

Viral quantification

Viral stocks and collected samples were titrated by tissue culture infectious dose 50% (TCID50/ml) in VeroE6 cells, using the Reed & Muench statistical method. In parallel, relative quantification of viral genome was performed by one-step real-time quantitative reverse transcriptase and polymerase chain reaction (RT-qPCR) from viral or total RNA extracted using QiAmp viral RNA or RNeasy Mini Kit (Qiagen) in the case of supernatants/apical washings or cell lysates, respectively. Primer and probe sequences (**Table S1**) were selected from those designed by the School of Public Health/University of Hong Kong (Leo Poon, Daniel Chu and Malik Peiris) and synthetized by Eurogentec. Real-time one-step RT-qPCR was performed using the EXPRESS One-Step Superscript™ qRT-PCR Kit (Invitrogen, reference 1178101K), in a 20 µl reaction volume containing 10 µl of Express qPCR supermix at 2X, 1 µl of forward primer at 10 µM, 1 µl of reverse primer at 10 µM, 0.5 µl of probe at 10 µM, 3.1 µl of PCR-water (Qiagen, reference 17000-10), 0.4 µl of Rox dye at 25 µM, and 2 µl of vRNA template. Thermal cycling was performed in a StepOnePlus™ Real-Time PCR System (Applied Biosystems) in MicroAmp™ Fast Optical 96-well reaction plates (Applied Biosystems, reference 4346907). Cycling conditions were as follows: reverse transcription at 50 °C during 15 min, followed by initial polymerase activation at 95 °C for 2 min, and then 40 cycles of denaturation at 95 °C for 15 sec and annealing/extension at 60 °C for 1 minute.

**Table S1. SARS-CoV-2-specific primer and probes used for viral genome quantification.**

| | |
|---|---|
| Target: ORF1b-nsp14 | |
| Forward primer (HKU-ORF1b-nsp14F) | 5'-TGGGGYTTTACRGGTAACCT-3' |
| Reverse primer (HKU- ORF1b-nsp14R) | 5'-AACRCGCTTAACAAAGCACTC-3' |
| Probe (HKU-ORF1b-nsp141P) | 5'-FAM-TAGTTGTGATGCWATCATGACTAG-TAMRA-3' |

Viral replication kinetics and antiviral treatment in VeroE6 cells

VeroE6 cells were seeded 24 h in advance in multi-well 6 plates, washed twice with PBS and then infected with SARS-CoV-2 at the indicated MOIs. For replication kinetics studies, supernatant samples were collected at different time-points were separated into 2 tubes: one for TCID50 viral titration and one RT-qPCR. For treatment studies, the inoculum of infected VeroE6 was removed 1 hpi and cells were immediately treated with solutions in DMEM of candidate molecules alone or in combination. DMEM alone (for diltiazem) or DMEM containing a DMSO concentration equivalent to that of the highest remdesivir dose tested was used as mock-treatment control. Supernatants were collected at 48 and 72 hpi and stored at -80 °C for RNA extraction and viral titration.

Viral infection and treatment in reconstituted human airway epithelia (HAE)

MucilAirTM HAE reconstituted from human primary cells obtained from nasal or bronchial biopsies, were provided by Epithelix SARL (Geneva, Switzerland) and maintained in air-liquid interphase with specific culture medium in Costar Transwell inserts (Corning, NY, USA) according to the manufacturer’s instructions. For infection experiments, apical poles were gently washed twice with warm OptiMEM medium (Gibco, ThermoFisher Scientific) and then infected directly with nasal swab samples or a 150 µl dilution of virus in OptiMEM medium, at a multiplicity of infection (MOI) of 0.1. For mock infection, the same procedure was performed using OptiMEM as inoculum. Samples collected from apical washes or basolateral medium at different time-points were separated into 2 tubes: one for TCID50 viral titration and one RT-qPCR. HAE cells were harvested in RLT buffer (Qiagen) and total ARN was extracted using the RNeasy Mini Kit (Qiagen) for subsequent RT-qPCR and Nanostring assays. Treatments with specific dilutions of candidate molecules alone or in combination in MucilAir® culture medium were applied through basolateral poles. All treatments were initiated on day 0 (5 1h after viral infection) and continued once daily at 24 and 48 hpi (2 and 3 treatments in total for samples collected at 48 and 72 hpi, respectively). Variations in transepithelial electrical resistance (Δ TEER) were measured using a dedicated volt-ohm meter (EVOM2, Epithelial Volt/Ohm Meter for TEER) and expressed as Ohm/cm2.

Transmission electron microscopy

Infected nasal and bronchial HAE were fixed with 2% glutaraldehyde (EMS) in 0.1 M sodium cacodylate (pH 7.4) buffer at room temperature for 30 min. After washing three times in 0.2 M sodium cacodylate buffer, cell cultures were post-fixed with 2% osmium tetroxide (EMS) at room temperature for 1 hour and deshydrated in a graded series of ethanol at room temperature and embedded in Epon. After polymerization, ultrathin sections (100 nm) were cut on a UCT (Leica) ultramicrotome and collected on 200 mesh grids. Sections were stained with uranyl acetate and lead citrate before observations on a Jeol 1400JEM (Tokyo, Japan) transmission electron microscope, equipped with an Orius 600 camera and Digital Micrograph.

Nanostring gene expression analysis

Briefly, after on-column mRNA extraction, gene expression levels was evaluated using two different gene panels (96 and 12 genes) using the Nanostring technology. Data processing and normalization were performed with nSolver analysis software (version 4.0, NanoString technologies) and results are expressed in fold change induction compared to the mock condition. Heatmap and Principal Component Analysis (PCA) were carried out using Genomics Suite 7 (Partek, St Louis, MO, USA).

**Fig. S1**

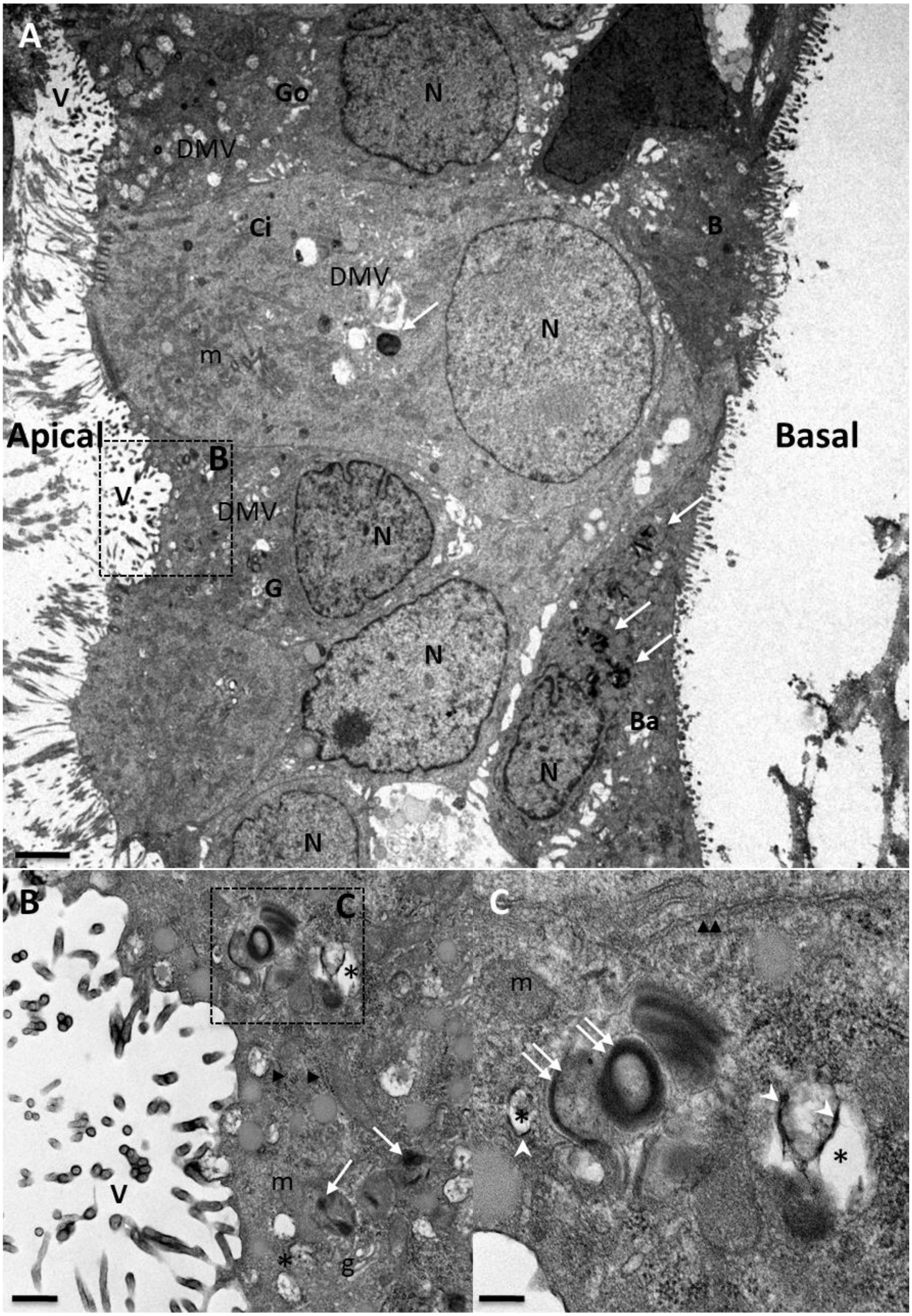

**Fig. S1. Electron micrographs of reconstituted nasal human airway epithelia (HAE) directly inoculated with SARS-CoV-2-positive clinical samples.** MucilAir™ nasal HAE were directly inoculated on the apical surface with a nasal swabs sample from a COVID-19 patient. Forty-eight hours post inoculation, HAE were fixed and processed for transmission electron microscopy analysis, as described in Materials and Methods. **(A)** Thin section cut through inoculated MucilAir™ showing ciliated cells (Ci), goblet cells (Go) and basal cells (Ba). Some typical features of coronavirus-induced cell ultrastructure remodeling and large electron-dense accumulation of viral materials (white arrow) are both observed in apical and basal cells. Apical and basal poles are indicated. Scale bar: 2 µm. **(B)** Enlargements of an infected mucus cell. Virion progeny is observed at the surface of mucus cells. Scale bar: 0.5 µm. **(C)** Detailed view of characteristic virus-induced single-membrane vesicles (asterisk) showing their electron-lucent interior with a spider web-like content and some pieces of double membrane (indicated by white arrowhead) and strong accumulations of viral nucleocapsids (double white arrow) indicating viral replication sites. Golgi vesicles (g), rough (arrowhead) and smooth (double arrowhead) endoplasmic reticulum are indicated. Scale bar: 2 µm. N: nucleus; DMV: cytoplasmic double-membrane vesicles; m: mitochondria.

**Fig. S2**

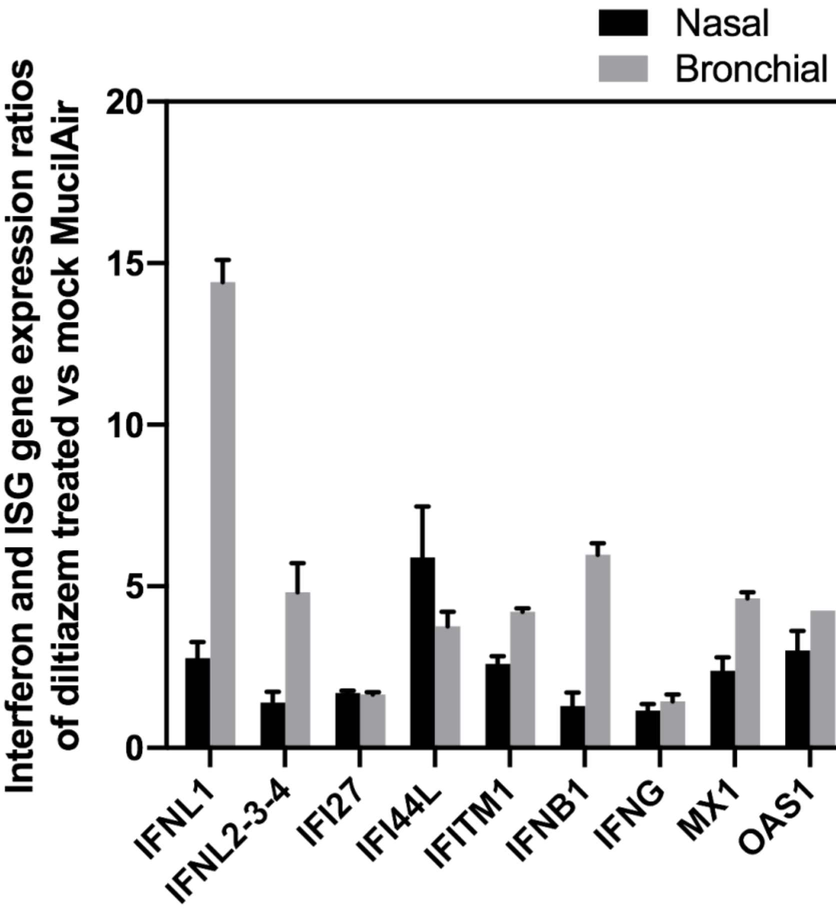


**Fig. S2. Diltiazem induces type III IFN-related gene expression in human airway epithelia (HAE).** The differential expression of type III IFNs (12-gene panel) was evaluated using the Nanostring technology at 72 hours post-treatment in diltiazem-treated nasal and bronchial HAE treated by diltiazem compared to mock-treated controls. Data treatment and normalization ware performed with nSolver analysis software (version 4.0, NanoString technologies). Results are expressed as mRNA expression ratios of IFNL1, IFNL2, 3, 4, IFNB1, IFNG and ISG (IFI27, IFI44L, IFITM1, Mx1, OAS1) genes compared to the mock condition. Representative data are shown from three independent experiments.

**Data S1. Innate immune transcriptional signature during the time course of SARS-COV-2 infection.** Briefly, after on-columns mRNA extraction, expression was evaluated using two different gene panels (composed of 96 and 12 genes respectively) using the Nanostring technology. Data treatment and normalization ware performed with nSolver analysis software (version 4.0, NanoString technologies). Results are expressed in counts and fold change induction compared to the mock condition.

(submitted as a separate .xlsx file)
